# Monocyte single cell-type gene expression measured in peripheral blood by DIRECT LS-TA method: the ratio-based biomarkers of (*IFI27/PSAP*) showed superior performance than interferon score in triage patients with viral infection

**DOI:** 10.1101/2025.05.08.652590

**Authors:** Nelson LS Tang, Tsz-Ki Kwan, Dan Huang, JH Huang, XY Wang, Grace CY Lui, Suk-Ling Ma, Kwong-Sak Leung

## Abstract

A rapid method to triage febrile patients into different categories of etiologies remains a significant challenge even nowadays, when many molecular tests for pathogens are available. Routine serum protein tests like C-reactive protein and procalcitonin have limited specificity. Host response gene signatures are promising biomarkers but they usually require assaying many genes, e.g. 7 genes are commonly used to calculate the interferon (IFN) score. However, these gene panels fail to capture cell-type-specific host responses. Measuring gene expression of a specified single cell population, like monocytes, offers enhanced biological insight. However, it currently requires laborious cell sorting or costly single-cell sequencing techniques, limiting its clinical applicability.

This study aims to develop a simple ratio-based biomarker (RBB) representing monocyte-specific host response to viral infection called DIRECT LS-TA method. A simple ratio of 2 genes (both are shortlist monocyte informative genes) quantified in peripheral blood (PB) samples correlated with gene expression in purified monocytes in the corresponding individual. These RBBs cover 3 interferon-stimulated genes (ISGs): *IFI27/PSAP*, *IFI44L/PSAP* and *SIGLEC1/PSAP*. They are compared to the conventional multi-gene IFN score in the differentiation of viral infection.

Public gene expression datasets from NCBI GEO were used to shortlist monocyte-informative genes that can be used as the RBB in PB. The DIRECT LS-TA RBB was calculated as the ratio of the target ISG transcript abundance (TA) to that of another reference gene (*PSAP* or *CTSS)* directly quantified from bulk PB data (e.g., Log(*IFI27/PSAP*) in WB). The correlation (expressed by coefficient of determination, R²) between these DIRECT LS-TA RBBs and the gold-standard target gene TA measured in purified monocytes was assessed. The diagnostic performance of selected RBBs (*IFI27/PSAP*, *IFI44L/PSAP*, *SIGLEC1/PSAP)* was compared against the conventional 8-gene IFN score for differentiating viral infections from controls.

Direct LS-TA RBBs measured in PB showed strong correlation with gold-standard gene expression measured in purified monocytes (R^2^ ranged from 0.53 for the target gene *IFI27* to >0.9 for the target gene *IFI44L*). This high level of correlation supports that this simple RBB (DIRECT LS-TA) method can replace the tedious cell sorting approach to obtain single-cell-type gene expression data. All DIRECT LS-TA results of ISGs were raised during viral infection. The best clinical performance in triaging viral infection patients was achieved by *IFI27/PSAP* or *IFI27/CTSS* across all datasets. For example, in the GSE111368 dataset, *IFI27/PSAP* achieved an AUC of 0.94 (95% CI 0.90-0.97) with 88% sensitivity and 95% specificity, surpassing the IFN score’s AUC of 0.90 (95% CI 0.85-0.94) with 79% sensitivity and 93% specificity.

**Conclusion:** The DIRECT LS-TA method, utilizing the format of simple two-gene ratio-based biomarkers like *IFI27/PSAP*, provides a robust and accurate measure of monocyte-specific interferon pathway activation directly from peripheral blood. The superior performance of the DIRECT LS-TA method makes it a promising, readily implementable tool for clinical triage. Its ability to provide single-cell-type specific information, rapid turnaround using standard qPCR/dPCR technology, and enhanced biological specificity make it a valuable molecular host response assessment.

## 1. Introduction

Febrile illness is a common clinical presentation, necessitating rapid and accurate triage to guide appropriate management. Current diagnostic approaches often rely on the detection of specific pathogens by culture or molecular methods which can be time-consuming and do not always identify the causative agent. While the host responses differently to different categories of pathogens (e.g. viral or bacterial), biomarkers targeting host response are less well developed. Currently, only few host response biomarkers are available and most of them are serum proteins, such as C-reactive protein (CRP) and procalcitonin (Leticia Fernandez-Carballo et al. 2021) . They can indicate the presence of inflammation, however they lack the specificity required to differentiate between various underlying etiologies.

### 1.1 Activation of interferon (IFN) stimulated genes during viral infection

Viral infection leads to activation of Type I interferons in infected epithelial cells by the virus. Interferon-stimulated genes (ISGs) play a fundamental role in the innate immune system, act as inhibitors of viral replication in infected cells, and have a defensive action in uninfected cells. Mommert-Tripon et al incorporated nasal IFN response into clinical diagnosis of viral infection (Mommert-Tripon et al. 2024). On the other hand, IFN response in blood is more accessible. The levels of expression of a battery of interferon-stimulated genes (ISGs) in peripheral blood (commonly known as interferon score, IFN score) were first developed as bioassay of systemic IFN responses in autoimmune diseases, e.g. SLE (Kim et al. 2018; Feng et al. 2006; Higgs et al. 2011). Typically, quantification of expression of 5 ISGs (e.g. *IFI44L, IFIT1, IFI27, SIGLEC1, IFITM3*) and 2-3 house-keeping genes (i.e. total 7-8 genes) are required to calculate the IFN score (Nocturne and Mariette 2022). Recently, the ratio of *IFI44L* / one house-keeping gene is proposed to be as useful as the multi-gene IFN score in the diagnosis of patients with interferonopathy (Adang et al. 2024). The authors reported that simple ratio-based biomarker (RBB) approach performed as good as the 7-gene IFN scores.

We and others studied host response in patients during the attack of severe acute respiratory syndrome (SARS) caused by the virus SARS-CoV-1 between 2002-2004 and found that serum protein IP-10 (Interferon gamma-induced protein 10) which is encoded by an ISG (*CXCL10*) was a risk factor for severe infection (Tang et al. 2005). So, activation of interferon pathway could serve as potential biomarkers in viral infection. However, most attempts up to now were carried out using serum proteins like interferon alpha and IP-10, rather than ISG. Trouillet-Assant et al compared interferon score and serum interferon-alpha protein level between patients with viral and bacterial infection and showed that they had comparable sensitivity (Trouillet-Assant et al. 2020) . Area under curves of both biomarkers were above 0.9 which was better than other biomarkers currently in clinical use (e.g. CPR). Similarly, during the COVID-19 epidemics, we and other teams also showed activation of ISG during infection which confirmed that interferon score was a potential biomarker for viral infection in general (Moradi Marjaneh et al. 2023; Huang et al. 2023).

### 1.2 Pitfalls of using DEG biomarkers in blood samples

The typical approach to discover gene expression biomarkers is through genome-wide gene expression analysis of peripheral blood of infected patients and healthy controls. Then genes showing different expression between the 2 groups are identified as differentially expressed genes (DEG). Tens to hundreds of these DEGs can be used as the gene signature of viral infection. Examples of these signatures include those developed by Li et al (Li et al.) , Rao et al (Rao et al.) and Xu et al (Xu et al. 2021). For example, a 29 genes panel of DEG was found useful to differentiate viral from bacterial infections (Bauer et al. 2021). Other example panels of over 30 genes have also been proposed (Bodkin et al. 2022; Tsalik et al. 2016; Ko et al. 2022). Some of them are also available as laboratory-developed tests (LDT) only provided by special laboratories(Holcomb et al. 2017). Recent study suggested that diagnostic performance was clinically acceptable and the area-under-curve (AUC) was around 0.89 for diagnosis of viral infection (Tong-Minh et al. 2025) . However, due to the large number of genes required to analyse, these tests are only available at special laboratories or require to use highly special equipment (Tsalik et al. 2016). It largely limits the availability of such assays in clinics or emergency department. Therefore, a simple gene expression biomarker that is readily implement on equipment commonly available in hospital laboratory is still not available and is a topic of active research.

Current approaches of using transcript abundance (TA) of DEG in peripheral blood samples as biomarkers are performed face a common unresolved fundamental problem. That is the cell mixture nature of peripheral blood. PB DEG biomarkers convey no information about the TA in any specific leukocyte population (or a single cell-type, such as monocytes or lymphocytes). Taking IFN score test as an example, it is not aware which cell-type produces mRNA of *IFI44L* or *IFIT1* genes. Any change in TA in PB could be caused by changes in the cell count proportion among leukocyte populations or a genuine activation of gene transcription in one or more cell populations. In order to obtain single cell-type TA, additional laboratory procedures are required to carry out. The conventional methods to obtain single cell-type TA include cell sorting by flow cytometry or magnetic beads which have been taken as the gold standard (e.g. quantification of TA after cell sorting to isolate monocytes from PB) (Quach et al. 2023). Recent developments in single cell RNA (scRNA) sequencing have make it possible to obtain the full genome expression profile for every individual cells. However, cell sorting and scRNA sequencing are either too labour intensive or costly to make them feasible for everyday clinical application in hospital laboratories.

### 1.3 A simple ratio-based biomarker could reflect single cell-type TA in PB

We recently described a direct method to determine TA of shortlisted genes predominantly produced by monocyte in PB using DIRECT LS-TA method. The DIRECT LS-TA method first shortlists single cell-type informative genes and then select 2 genes among them to form a RBB (with a target gene and another denominator gene that are specific for the cell-type of interest) (Tang et al. 2025). This RBB can estimate single cell-type specific gene expression from bulk blood samples without the need for complex cell separation techniques. These RBBs have been shown by the high correlation between the gold-standard TA determined in purified samples of the cell-type of interest (e.g. monocytes). Such simple ratio-based biomarker can be readily implementation in hospital laboratories as it is easy to standardise and no special machine is required.

Among those shortlisted genes predominantly expressed by monocytes in PB are several ISGs (Tang et al. 2025). These ISGs include *IFI44L, SIGLEC1, and CXCL10*. Here, we also show in addition that IFI27 is also predominantly expressed by monocytes in PB. We hypothesize that RBBs based on these ISGs can both reflect their expression in monocytes and effectively differentiate viral infections from other febrile conditions. We selected *IFI27, IFI44L* and *SIGLEC1*, all well-established ISGs, and paired them with PSAP, a monocyte-specific reference gene, to form the RBBs which are (*IFI27/PSAP*), (*IFI44L/PSAP*) and (*SIGLEC1/PSAP*). Their diagnostic performance was compared to that of conventional IFN score in differentiation of viral infection. Through this approach, we aim to develop a simple, rapid, and cost-effective cell-type specific assays directly applied to PB samples for the triage of febrile illness.

## 2. Materials and methods

### 2.1 Datasets Used in the Analysis of Gene Expression of Peripheral Blood and Monocytes to identify ISG predominantly expressed by monocytes in PB

Similar to our previous report (Tang et al. 2025) , gene expression datasets obtained from peripheral blood samples were collected from the Gene Expression Omnibus (GEO), maintained by the US National Institutes of Health. The types of peripheral blood samples included whole blood (WB) and peripheral blood mononuclear cells (PBMCs). Specific cell types that have been further isolated and purified, such as isolated and purified monocytes, were also included in some datasets.

### 2.2 Datasets used in comparison of DIRECT LS-TA and IFN score in differentiation of viral infection

Only datasets with large collection of PB samples from viral infection patients (at least 40 viral infected patients) were evaluated. A main drawback of previous studies was the use of many datasets with small number of patients, such as datasets with even less than 10 patients. First, these datasets may not contribute much to the gene signal. Even worse, they added to exacerbate the noise in the overall analysis, e.g. between-batch and between-platform errors to the overall analysis. These confounding effects may lead to the identification of false positive biomarkers.

### 2.3 DIRECT LS-TA method shortlists Monocyte Informative Genes Whose Expression are predominantly from monocytes in cell-mixture Samples e.g. PBMC, WB

In our previous publication, we defined cell-type informative genes as genes which are predominantly expressed by only one single cell-type (e.g. monocyte) to the extent that ≥50% of gene transcripts of these informative genes in the cell-mixture sample (for example, PBMCs) were contributed by that single cell-type (Huang et al. 2021). Typically, the cell count percentage of monocytes in PBMCs was 10%-30% (Meskini et al. 2024; Nielsen et al. 2020). The proportional cell count of monocytes in PBMCs was pre-defined to be 20% in this case. By using the definition we described previously (Huang et al. 2021), assuming the proportional cell count of monocyte is 20%, the expression of an informative gene in the purified monocyte sample needed to be 2.5 times higher than that in the cell-mixture sample. This principle of this calculation is illustrated in Figure 1. The monocyte informative genes in the cell-mixture blood sample were shortlisted by using these conditions. Expression data from the isolated monocyte sample and cell-mixture sample (PBMCs or WB) in datasets from GEO (Table 1) were used to determine which genes fulfilled this requirement for being potential monocyte informative genes.

**Figure 1.**
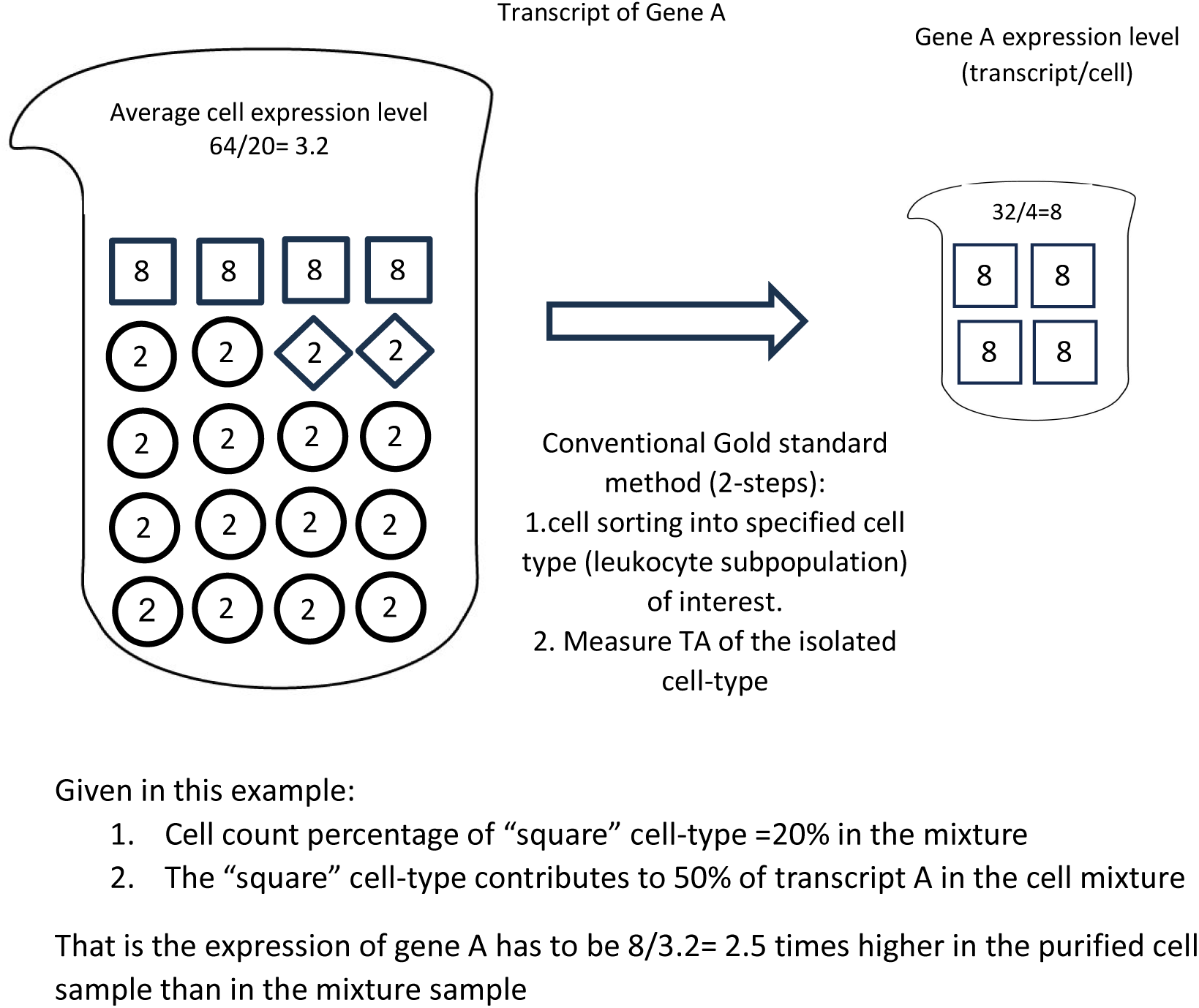
A schematic diagram of screening for cell subpopulation informative genes. The figure shows 2 samples, on left side is the original cell-mixture samples (e.g. PB) and 3 constitutional cell-types samples obtained from the original cell by cell separation / sorting, on the right side. Monocyte, the cell-type of interest, is shown as cells in square symbols. A pre-defined cell count percentage of monocyte is set to 20% as in PBMC. If any gene, such as gene A, has an average cellular expression level in an isolated monocyte sample that is above 2.5x (folds) higher than its average cellular expression level in a cell-mixture sample (for the square cell subpopulation, 8 (for isolated square cells) / 3.2 (in cell-mixture sample) = 2.5 folds), this cell subpopulation (i.e., the square cell subpopulation) contributes 50% of gene A transcripts in the cell-mixture sample. The gene expression level is represented by a ratio of number of transcripts to the number of cells. For example, the cell-mixture sample has a total of 64 transcripts of Gene A in 20 cells, the gene expression value = 64 / 20 = 3.2 transcript/cell. Such gene expression values are similar to the concept of relative expression quantification using housekeeping gene to normalize target gene expression. Genes with expression level 2.5x (folds) higher in the isolated square cell subpopulation than in the cell-mixture sample are potential monocyte informative genes. In this example, to demonstrate the principle, it is assumed that the gene expression levels of other cells are known. In fact, we only need to know the expression levels of the specified cell sample after isolation and the corresponding cell mixture sample to identify the single cell-type informative gene.

**Table 1.** List of PBMC or WB gene expression datasets used to identify ISG predominantly expressed by monocyte.

| Dataset accession number | Usage | Type of peripheral blood sample (WB or PBMC) | References |
| --- | --- | --- | --- |
| GSE138746 | 1. To calculate the 90th percentile value of the fold of expression in monocytes vs. PBMCs, and find out genes that meet the criteria for monocyte informative gene;<br>2. To examine and determine the correlation between the biomarker of gene expression obtained by using Direct monocyte LS-TA and the gene expression of isolated and purified monocytes (the gold standard) | PBMC and isolated and purified monocytes | (Tao et al. 2021) |
| GSE114407 | Ditto | PBMC and isolated and purified monocytes | (Kuan et al. 2019) |
| GSE60424 | Ditto | Whole blood and isolated and purified monocytes | (Linsley et al. 2014) |
| GSE11755 | Ditto | Whole blood and isolated and purified monocytes, lymphocytes and granulocytes | NA |
| GSE19491 | Ditto | Whole blood and isolated and purified monocytes and lymphocytes | (Berry et al. 2010) |
| GSE107011 | To compare the gene expression of granulocytes and monocytes | PBMC and isolated and purified monocytes and granulocytes | (Monaco et al. 2019) |

**Table 2.**
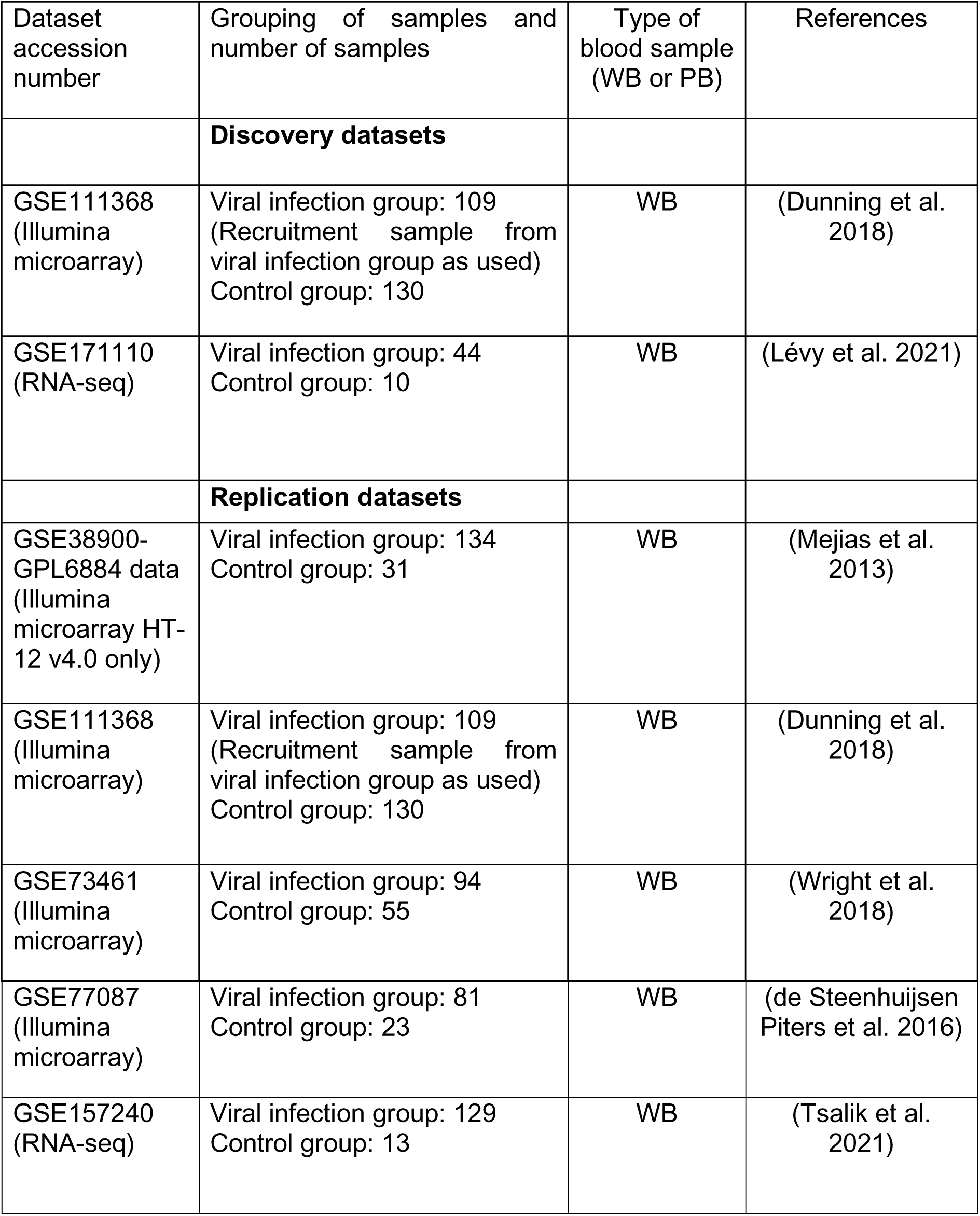
List of WB gene expression datasets used to in comparison of DIRECT LS-TA and IFN score in differentiation of viral infection.

### 2.4 RBBs using the Direct LS-TA method represent ISGs predominantly produced by monocytes in PB

Given a cell proportion of 20% for monocytes in PBMC, several known ISGs are shortlisted as monocyte informative genes as they fulfilled the requirement of having 2.5 times higher expression in purified monocytes than the corresponding PB sample collected from the same individual. This level of expression difference suggests that mRNA transcripts in PB are predominantly (>50%) produced by monocytes. Examples of ISGs that fulfilled monocyte informative gene criteria include *IFI27, IFI44L*, *SIGLEC1* and other listed in Table 3.

**Table 3.** List of Monocyte cell-type specific Informative Target Genes and 2 Reference Genes (*CTSS* and *PSAP*) * *CTSS* and *PSAP* are used as denominator genes in the RBB, so they do not have values in the column of correlation results when compare with the gold standard method. #*IFI27* results are further explained in the text.

| Monocyte Informative Reference and Target genes | The 90<sup>th</sup> percentile value of the fold of expression in monocytes vs. peripheral blood or PBMCs | Coefficient of variation (CV%) | Correlation between the Monocyte DIRECT LS-TA biomarker obtained directly from peripheral blood samples and the gold standard (expression level of target genes in isolated and purified monocytes) (coefficient of determination, R<sup>2</sup>) | Database source showing the best correlation |
| --- | --- | --- | --- | --- |
| Reference genes |  |  |  |  |
| *CTSS* | 2.8x | 9% | NA* |  |
| *PSAP* | 3.2x | 11% | NA* |  |
| Target genes |  |  |  |  |
| *CXCL10* | 3.6x | 212% | 0.84 | GSE60424 |
| *IFI27* | 9.1x | 130% | 0.92<sup>#</sup> | GSE11755 |
| *IFI30* | 3.3x | 20% | 0.60 | GSE60424 |
| *IFI44L* | 3.2x | 141% | 0.95 | GSE60424 |
| *IFITM3* | 2.7x | 100% | 0.87 | GSE138746 |
| *SIGLEC1* | 3.8x | 171% | 0.94 | GSE138746 |

In our previous study of COVID patients, we found that *IFI27* were also highly produced by monocytes (Huang et al. 2023). However, the correlation between Monocyte DIRECT LS-TA (*IFI27/PSAP*) biomarker obtained directly from peripheral blood samples and the gold standard (*IFI27/ACTB*) expression in purified monocytes were low in dataset GSE138746, R^2^=0.34. Although the correlation was statistically significant, R^2^ was below 0.5. Therefore, two additional datasets with both purified monocyte and PB samples were analysed (GSE19491 and GSE11755). Both datasets contained patients’ data with more intense *IFI27* activation in monocytes, while other datasets in Table 1 were collected mostly from healthy controls who had very low *IFI27* expression at the baseline levels. In order to exclude samples with very low *IFI27*, a filter was applied to include only patients with activated *IFI27* level in monocytes which was defined by a log ratio of *IFI27* to a housekeeping gene ACTB (beta-actin) of greater than - 8. And then the correlation between *IFI27* expression in purified monocytes and PB (DIRECT LS-TA) was re-evaluated (Table 4).

**Table 4.** The correlation between DIRECT LS-TA results, RBBs calculated from PB vs the actual gene expression level in purified monocytes.

| Dataset | R<sup>2</sup> when All samples included | R<sup>2</sup> when Only high *IFI27* expression samples (log (*IFI27/ACTB*) >-8 in purified monocyte) | Number of data points after filtering |
| --- | --- | --- | --- |
| GSE19491 | 0.53 (p=0.011) | 0.53 (p=0.011) | 11 |
| GSE11755 | 0.92 (p=0.00014) | 0.92 (p=0.00014) | 8 |
| GSE138746 | 0.34 (p<1e-7) | 0.81(p=0.0058) | 7 |
| GSE114407 | 0.31 (p=0.0002) | R<sup>2</sup> not determined | Only 3 data points |
| GSE60424 | 0.09 (p=0.22) | R<sup>2</sup> not determined | Only 3 data points |

### 2.5 Determining DIRECT LS-TA (RBB) results from gene expression datasets and conversion to multiple of median (MoM)

To develop the ratio-based biomarker (RBB), *PSAP* and *CTSS* which had the least biological variation were selected as the denominator genes among the shortlisted monocyte informative genes. These denominator gene are called monocyte informative reference gene which are selected based on having the least between-individual variance. Therefore, the percentage coefficient of variation (CV%) was calculated for each monocyte informative gene to find those with lowest CV%. Conventional housekeeping genes were not useful as they are expressed across all cell-types in peripheral blood but not specific to monocyte. To obtain a single cell-type informative RBB, both numerator and the denominator genes have to be predominantly produced by the cell-type of interest in a cell-mixture sample.

#### 2.5.1 Calculation of DIRECT LS-TA results

The DIRECT LS-TA results are calculated as an RBB using the following formula:

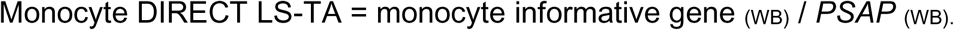

Or

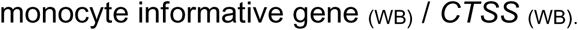

-depending on which denominator (PSAP or CTSS) is preferred, only one denominator is required in real application.

After log transformation, DIRECT LS-TA results can be expressed as:

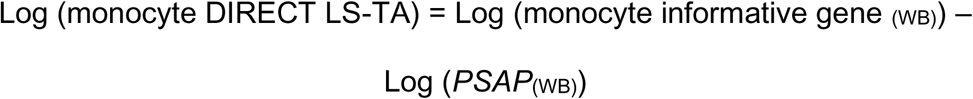

Or

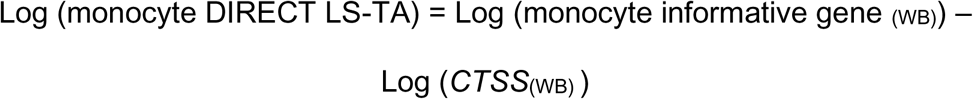

#### 2.5.2 MoM standardised fold changes of ISGs to compare across different studies and different TA assay platforms

When results from different assay platforms are analysed, the results of TA were expressed in very different units. One approach to compare across assay platforms is to convert DIRECT LS-TA results to fold-changes against healthy reference individuals. The standardised fold-change is called multiples of median (MoM), which can be calculated by subtracting DIRECT LS-TA results of patients by the median value in the control group. After log-transformation, MoM standardisation of individual results in effect assigned the median of log Monocyte DIRECT LS-TA of the control (healthy) group to zero. Then, the MoM value of log DIRECT LS-TA of individual sample is similar to the fold-change expressed as delta-delta CT values in qPCR experiments.

For comparison between DIRECT LS-TA RBB of ISGs and IFN score, we calculated the conventional IFN score using the following formula:

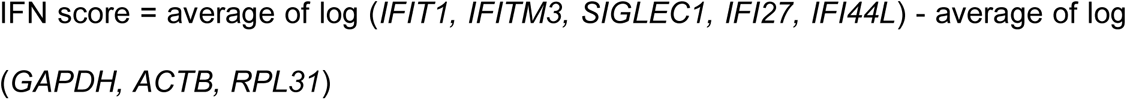

IFN score = average of log (*IFIT1, IFITM3, SIGLEC1, IFI27, IFI44L*) - average of log (*GAPDH, ACTB, RPL31*)

### 2.6 Statistical Analysis

Statistical analyses were performed using the R statistical package (R4.2.1). Associations between DIRECT LS-TA and categorical clinical variables were tested by Wilcoxon Rank sum test (two categories). For RNA-seq data, gene counts were normalized using the TPM method. The correlations between DIRECT LS-TA and each of the single genes were performed by Pearson correlation by R^2^.

### 2.7 Evaluation of Biomarker Performance

Six DIRECT LS-TA RBB (*IFI44L/PSAP*, *IFI44L/CTSS*, S*IGLEC1/PSAP*, *SIGLEC1/CTSS, IFI27/PSAP* and *IFI27/CTSS*) were extensively evaluated for their ability to differentiate viral infection from healthy controls. Area under the receiver operating characteristic curve (AUC-ROC) analysis was performed to assess diagnostic performance. Additionally, sensitivity and specificity at various cutoff points were calculated to determine optimal diagnostic thresholds. Receiver operating characteristic (ROC) curve analysis was performed to evaluate the diagnostic performance of each biomarker. Area under the curve (AUC), sensitivity, and specificity were calculated for:

1. DIRECT LS-TA *IFI27/PSAP* or *IFI27/CTSS*
2. DIRECT LS-TA *IFI44L/PSAP* or *IFI44L/CTSS*
3. DIRECT LS-TA *SIGLEC1/PSAP* or *SIGLEC1/CTSS*
4. Conventional IFN score (see formulae in the last section)

## 3. Results

### 3.1 Correlation between DIRECT LS-TA and Monocyte-specific Expression

Initial validation studies demonstrated that DIRECT LS-TA RBB results obtained from PB strongly correlated with gene expression measured in purified monocytes (Table 3). The correlation was particularly strong for *IFI44L/PSAP* and *IFI44L/CTSS* (both R^2^>0.9, p-value < 1e-8), confirming that this RBB method effectively captures monocyte-specific gene expression without the need for cell separation. Similar strong correlations were observed for *SIGLEC1/PSAP* and *CXCL10/PSAP* ratios, supporting the validity of using these DIRECT LS-TA measurements as proxies for monocyte-specific interferon responses. But the coefficient of determination (R^2^) between *IFI27/PSAP* and *IFI27* expression (*IFI27/ACTB*) in purified monocytes was lower than 0.5.

After filtering out monocyte samples with the very low baseline expression of log(*IFI27/ACTB*) below -8, the correlation between the *IFI27* expression in purified monocyte. In Figure 2, gold standard values of gene expression in purified monocytes are shown on the X-axis, and DIRECT LS-TA RBB are shown on the Y-axis. After removing very low expression samples, DIRECT LS-TA RBB results of *IFI27* (*IFI27/PSAP* in PB) showed much improved correlation with the ground-truth (Table 4).

**Figure 2.**
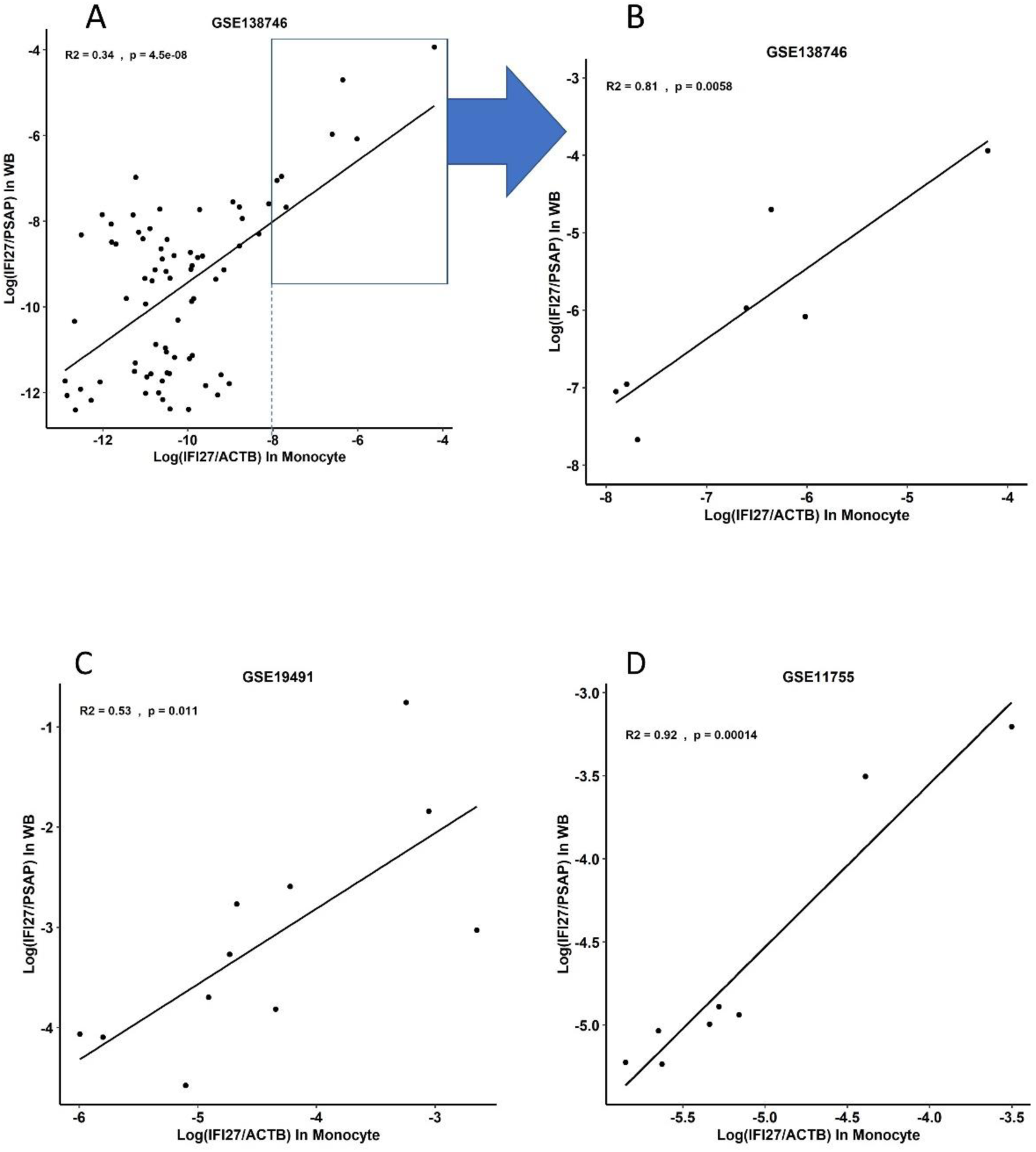
Correlation of DIRECT LS-TA of *IFI27* in PB with *IFI27* gene expression in purified monocyte (gold standard). Figure 2A and 2B are scatter plot showing DIRECT LS-TA results of *IFI27* in GSE138746 and many subjects had very low *IFI27* expression (such that log (*IFI27/ACTB* in purified monocyte) < -8). After confining the correlation analysis to those individuals with monocyte *IFI27* expression above that level (those inside the rectangle in Figure 2A), DIRECT LS-TA of *IFI27* was significantly correlated with *IFI27* expression in purified monocytes (R^2^=0.81, p=0.0058). Figure 2C and 2D show the correlation between DIRECT LS-TA of *IFI27* in PB and *IFI27* gene expression in purified monocyte in two other datasets.

### 3.2 Differential Expression Analysis in Viral Infection (Table 5)

Compared to healthy controls, we showed that DIRECT LS-TA values were significantly increased during viral infection across both discovery datasets (GSE111368 and GSE171110) (Figure 3). The most pronounced difference was observed for DIRECT LS-TA *IFI27,* which showed a highly significant elevation in viral infection patients (p<1e-9). Similarly, DIRECT LS-TA *IFI44L* and *SIGLEC1* were also significantly increased in the viral infection group (p<1e-5). These findings were consistent across both microarray and RNA-sequencing platforms, demonstrating the robustness of these biomarkers across different quantification methods.

**Figure 3.**
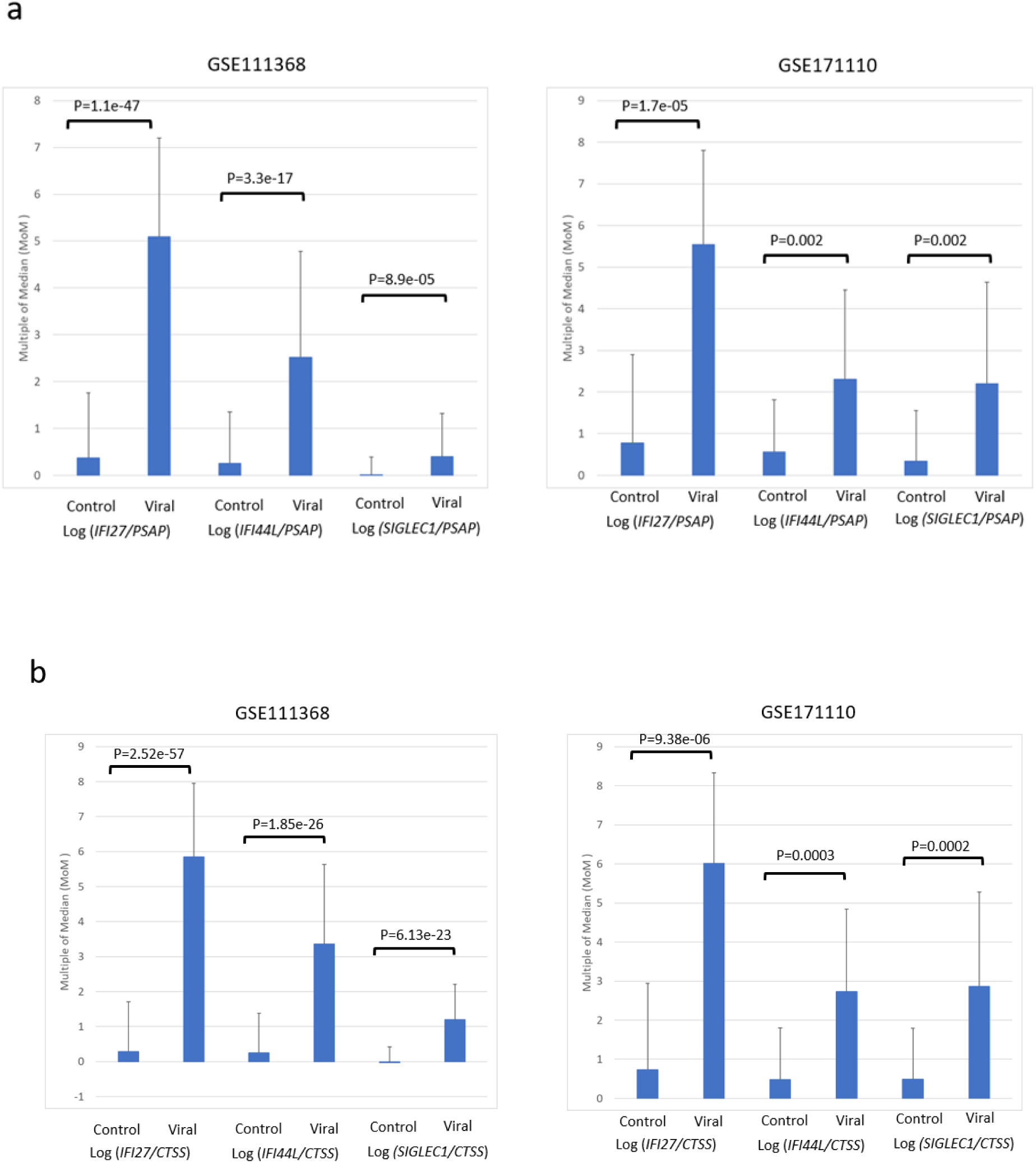
DIRECT LS-TA result in patients with viral infection compared to healthy control in GSE111368 and GSE171110 using (a) *PSAP* and (b) *CTSS* as reference gene. P value of student t-test was shown.

**Table 5.** DIRECT LS-TA results in patients with viral infection compared to healthy controls.

|  |  | Multiples of Median (MoM) values<br>mean, SD, (-Log<sub>10</sub>(P value of t-test)) |  |  |  |  |  |
| --- | --- | --- | --- | --- | --- | --- | --- |
| DIRECT LS-TA | Dataset | GSE111368 | GSE171110 | GSE38900-GPL6884 | GSE73461 | GSE77087 | GSE157240 |
| Log (*IFI27/PSAP*) | Control | 0.37,1.39 | 0.78,2.11 | 0.32,1.37 | 0.76,1.99 | 0.79,2.3 | 0.69,1.81 |
|  | Viral | 5.09,2.11,(46.97) | 5.55,2.25,(4.76) | 4.05,1.64,(17.46) | 5.61,2.20,(25.92) | 5.18,1.29,(8.52) | 6.02,2.86,(7.87) |
| Log (*IFI27/CTSS*) | Control | 0.29,1.42 | 0.73,2.20 | 0.35,1.44 | 0.46,2.02 | 0.90,2.43 | 0.71,1.87 |
|  | Viral | 5.85,2.10,(56.60) | 6.01,2.32,(5.03) | 4.13,1.73,(16.80) | 5.66,2.41,(27.24) | 5.68,1.47,(8.82) | 6.36,2.98,(8.05) |
| Log<br>(*IFI44L/PSAP*) | Control | 0.25,1.10 | 0.56,1.25 | 0.15,0.62 | 0.20,1.47 | 0.39,1.46 | 0.23,1.26 |
|  | Viral | 2.52,2.26,(16.48) | 2.3,2.15,(2.61) | 1.36,1.24,(11.16) | 2.59,1.65,(14.82) | 2.43,1.23,(6.08) | 3.50,1.90,(6.98) |
| Log<br>(*IFI44L/CTSS*) | Control | 0.25,1.13 | 0.48,1.32 | 0.20,0.66 | 0.40,1.41 | 0.31,1.56 | 0.25,1.31 |
|  | Viral | 3.36,2.27,(25.73) | 2.73,2.11,(3.48) | 1.47,1.27,(11.02) | 3.14,1.77,(18.25) | 2.74,1.32,(6.95) | 3.85,1.80,(7.21) |
| Log<br>(*SIGLEC1/PSAP*) | Control | 0.01,0.38 | 0.33,1.22 | 0.00,0.11 | -0.04,0.47 | 0.09,0.32 | 0.40,1.30 |
|  | Viral | 0.39,0.93,(4.05) | 2.2,2.44,(2.81) | 0.54,0.76,(12.57) | 0.57,0.75,(8.18) | 0.22,0.37,(1.01) | 3.44,1.99,(6.30) |
| Log<br>(*SIGLEC1/CTSS*) | Control | -0.02,0.44 | 0.49,1.29 | 0.01,0.15 | 0.06,0.70 | 0.00,0.42 | 0.31,1.35 |
|  | Viral | 1.20,1.00,(22.21) | 2.87,2.41,(3.70) | 0.61,0.84,(11.94) | 1.02,1.02,(9.69) | 0.53,0.52,(5.09) | 3.68,2.00,(6.67) |

The conventional IFN score, calculated as the average of five ISGs *(IFIT1, IFITM3, SIGLEC1, IFI27, IFI44L*) normalised to the mean of three housekeeping genes *(GAPDH, ACTB, RPL31*), was also significantly elevated in viral infection patients (p<1e-9). However, this method required quantification of eight genes compared to the two genes needed for DIRECT LS-TA measurements.

### 3.3 Comparative Analysis of Diagnostic Performance

To evaluate the clinical utility of these biomarkers, we performed detailed ROC curve analyses. In dataset GSE111368, DIRECT LS-TA *IFI27/PSAP* demonstrated superior diagnostic performance with an AUC of 0.94, surpassing both other DIRECT LS-TA measurements and the conventional IFN score (Table 6, Figure 4). The sensitivity and specificity for DIRECT LS-TA *IFI27* were 0.88 and 0.95, respectively, indicating excellent discriminatory power for viral infection.

**Figure 4.**
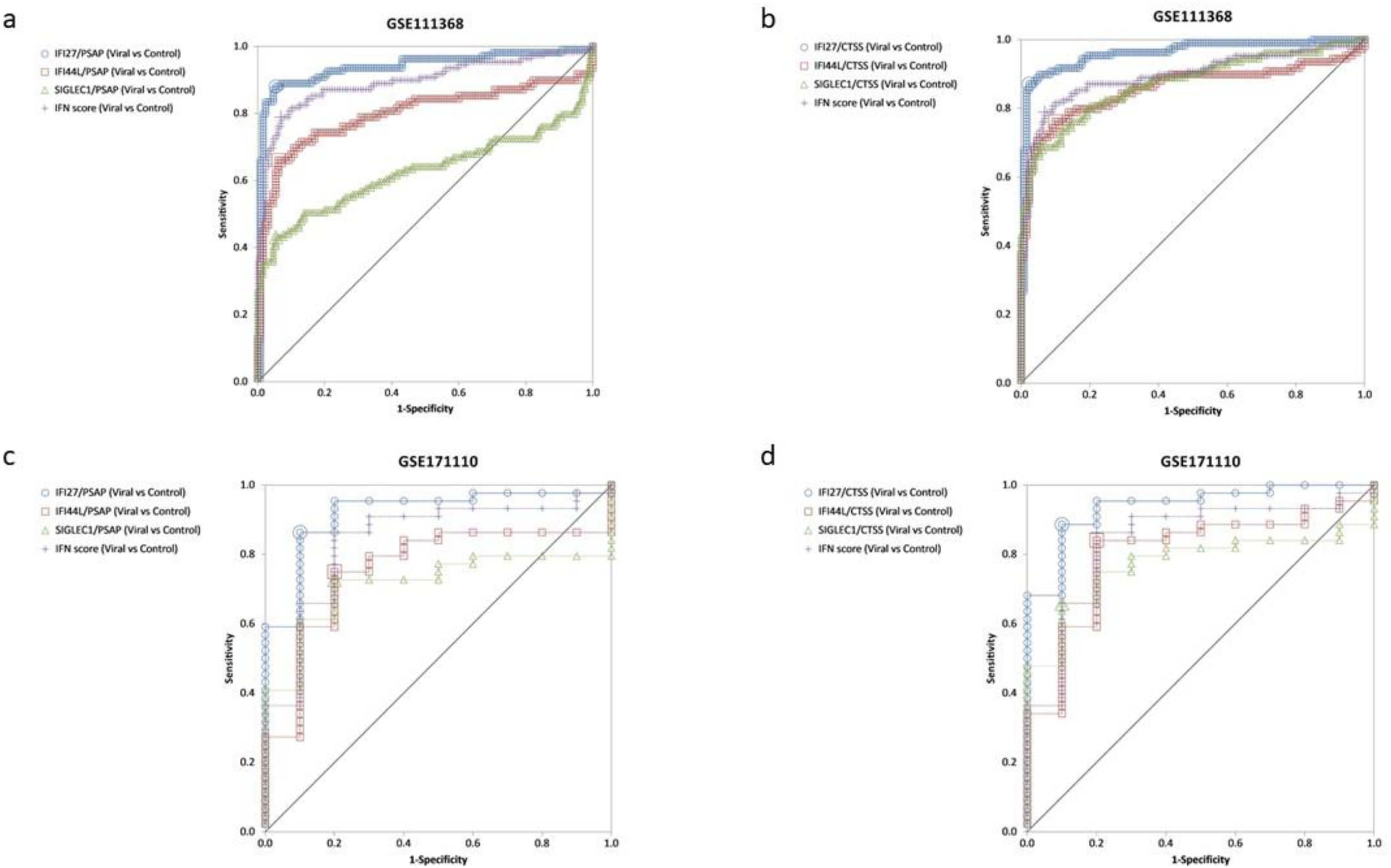
Receiver operating characteristic (ROC) curve of biomarkers in GSE111368 (a and b) and GSE171110 (c and d)

**Table 6.** Performance metrics of different biomarkers in differentiating viral infection.

| Biomarker | AUC ,95% confidence interval |  | Sensitivity, 95% confidence interval |  | Specificity, 95% confidence interval |  |
| --- | --- | --- | --- | --- | --- | --- |
|  | GSE111368 | GSE171110 | GSE111368 | GSE171110 | GSE111368 | GSE171110 |
| DIRECT LS-TA<br>*IFI27/PSAP* | 0.94,<br>0.90-0.97 | 0.92,<br>0.83-1 | 0.88,<br>0.80-0.93 | 0.86,<br>0.73-0.95 | 0.95,<br>0.89-0.98 | 0.9,<br>0.55-1 |
| DIRECT LS-TA<br>*IFI27/CTSS* | 0.96,<br>0.94-0.99 | 0.94,<br>0.87-1 | 0.87,<br>0.79-0.93 | 0.89,<br>0.75-0.96 | 0.98,<br>0.93-0.99 | 0.9,<br>0.55-1 |
| DIRECT LS-TA<br>*IFI44L/PSAP* | 0.80,<br>0.74-0.86 | 0.76,<br>0.60-0.91 | 0.66,<br>0.56- 0.75 | 0.75,<br>0.60- 0.87 | 0.94,<br>0.88-0.97 | 0.8,<br>0.44- 0.97 |
| DIRECT LS-TA<br>*IFI44L/CTSS* | 0.86,<br>0.80-0.91 | 0.80,<br>0.66-0.95 | 0.76,<br>0.67- 0.84 | 0.84,<br>0.70- 0.93 | 0.90,<br>0.84-0.95 | 0.8,<br>0.44- 0.97 |
| DIRECT LS-TA<br>*SIGLEC1/PSAP* | 0.63,<br>0.55-0.71 | 0.72,<br>0.58- 0.85 | 0.43,<br>0.34-0.53 | 0.73,<br>0.57- 0.85 | 0.95,<br>0.89-0.97 | 0.8,<br>0.44-0.97 |
| DIRECT LS-TA<br>*SIGLEC1/CTSS* | 0.87,<br>0.83-0.92 | 0.77,<br>0.64- 0.90 | 0.69,<br>0.59-0.77 | 0.66,<br>0.50- 0.79 | 0.95,<br>0.89-0.97 | 0.9,<br>0.55-0.99 |
| Conventional IFN<br>score | 0.90,<br>0.85-0.94 | 0.84,<br>0.70-0.98 | 0.79,<br>0.70-0.86 | 0.86,<br>0.73-0.95 | 0.93,<br>0.87-0.97 | 0.8,<br>0.44-0.97 |

The conventional IFN score showed good but slightly lower performance metrics, with an AUC of 0.9, sensitivity of 0.79, and specificity of 0.93. DIRECT LS-TA *IFI44L* and *SIGLEC1* showed moderate performance with AUCs of 0.80 and 0.63, respectively. The complete performance metrics for all biomarkers in both datasets GSE111368 and GSE171110 was shown in table 6 and receiver operating characteristic (ROC) curves of biomarkers in both datasets were shown in figure 4.

These comprehensive analyses demonstrate that DIRECT LS-TA monocyte *IFI27*, a simple two-gene ratio-based biomarker, outperforms the more complex conventional IFN score in identifying viral infections. The superior performance of this simplified approach, combined with its technical robustness and temporal stability, suggests that it could be a valuable tool for rapid diagnosis of viral infections in clinical settings.

## 4. Discussion

Biomarkers of host response to infection have great potential for clinical applications in triage of patients with fever which helps to delineate the type of pathogen causing the infection. Such differentiation is important as patients can be managed promptly with necessary samples are collected for investigation and therapy are given in time. Currently, only two serum proteins reflecting the host response, C-reactive protein (CRP) and procalcitonin (PCT) are in routine clinical use. However, these serum protein biomarkers do not convey any cell-type specific host response information, but just provides a crude representation of the overall systemic host response to infection.

With the advance in molecular techniques to quantify gene expression, many researchers analysed the TA in peripheral blood by microarray RNA-sequencing or on specific platforms (Mommert et al. 2021) . In the commonly practiced algorithms, the expression of each gene is statistically analysed one by one, and then the genes with the greatest expression difference between different groups were identified as the biomarker. These gene expression biomarkers are also called differential expression genes (DEGs). However, this method ignores the confounding factor of the cell counts of various cell subpopulations and their variations in different diseases. Therefore, variations in these factors will weaken the effectiveness of DEG biomarkers in differentiating diseases.

Peripheral blood is composed of various cell-types including various leukocyte subpopulations such as monocyte, lymphocyte and granulocyte which are present in different proportions. The gene expression level in a single leukocyte cell-type (e.g. monocytes) in blood is highly promising as an informative biomarker. But single leukocyte cell-type transcript abundance is not readily measured by available methods in the past. Currently, measurement of change in transcript abundance (TA) in blood samples in most studies are DEGs which do not quantify TA in a specific single leukocyte cell-type. Any change in gene expression of DEGs could represent change in TA of one/more leukocyte cell-types, change in cell count proportions of one or more cell-types or both. Here, we described a simple method (DIRECT LS-TA) to detect the change in gene expression in a single leukocyte cell-type, monocytes by a RBB (e.g. IFI27/PSPA) quantified directly in peripheral blood samples. It avoids both the labour-intensive procedure of cell isolation and high cost of single cell sequencing. Furthermore, DIRECT LS-TA has enhanced specificity over DEG which will make it a more reproducible biomarker as it reflects biology of the underlying host response in a single leukocyte population.

Computation algorithms have developed to deconvolute the cell-count proportion of each cell type presented in a peripheral blood sample using matrix deconvolution. However, these algorithms assumed the same expression profile for each cell-types for all subjects in a group (Shen-Orr et al. 2010). Only recently, methods are developed to determine single cell-type gene expression for individual subjects in a dataset using machine learning methods (Khatri et al. 2024) . However, all these methods require input of the gene expression data of the whole genome such as microarray data or RNA-sequencing data. And these technologies are not ready for everyday clinical use at the moment or still too expensive to apply. Of course, the gold standard approach is to isolate monocyte from peripheral blood and measure gene expression of the target genes. However, it requires cell separation which is the cumbersome and technically challenging in a clinical laboratory setting. In fact, it is not practical to implement cell isolation procedures in routine hospital laboratory for the time being.

By using DIRECT LS-TA method, gene expression of a single cell-type in peripheral blood can be directly determined for every single sample which makes it a valuable technology for translation to clinical use. The correlation of DIRECT LS-TA results and TA in isolated monocytes was very strong, R^2^ for some target genes (e.g. *IFI44L*) were up to 0.9 or even more. DIRECT LS-TA results represent the average gene expression of a single leukocyte cell-type, and therefore, is not confounded by the cell count proportions in cell-mixture samples. Furthermore, this ratio-based biomarker method can be readily translational to clinical application as it only requires the use of qPCR or dPCR machines which are widely available nowadays in most clinical laboratories.

*IFI27/PSAP* turned out to be the champion in differentiation of viral disease in febrile patients. This RBB represents monocyte single cell-type expression of the interferon induced gene, *IFI27*. *IFI27* has low expression in PB / monocyte at baseline but it is highly expressed after IFN stimulation. Therefore, in the 2 studies with infection patients, DIRECT LS-TA of *IFI27* (i.e. *IFI27/PSAP*) in PB and *IFI27* in purified monocytes showed good correlations with R^2^ between 0.53 and 0.92. Further examination of the purified monocyte expression of *IFI27* showed that many patients had activated *IFI27* which was 2-3 times above baseline *IFI27*. So we focused on those patients with IFN activation and re-evaluated the correlation. The enhanced correlation after exclusion of the baseline samples support that *IFI27* is a monocyte informative gene when it is activated.

Like innate immunity towards other virus, the ISGs activation was observed largely during the early course of infection (before day 10). As sampling time and age were two major confounders of ISG expression, they may account for contradicting observations among previous studies. In the future application, DIRECT LS-TA monocyte should be performed when the febrile patient is waiting at the ER or early after admission.

Results show that DIRECT LS-TA for *IFI27* (equal to the ratio of *IFI27* to *PSAP*) in whole blood was the best indicator of viral infection. This 2-gene ratio-based biomarker performed even better than the conventional IFN score which required quantification of 8 genes. This finding is consistent with our previous study of IFN score in COVID patients where a small percentage of patients had “normal” IFN score but only *IFI27* remained elevated to a small extent while other IFN-stimulated genes returned to normal level (Huang et al. 2023).

The important role of monocytes in the activation of these ISG after a viral infection is also confirmed by using scRNA-seq data. It is clear that these genes were preferentially activated in the monocyte population, particularly after infection. Therefore, using the DIRECT LS-TA results as a single cell-type specific ratio-based biomarker is advantageous over using bulk gene expression results in patient triage.

### Limitations

Our study was limited by confining to use publicly available gene expression datasets and having little control of the design of the original study e.g. case definition of viral infection and different platforms of gene expression quantification. Moreover, our study and results could only be used to discriminate viral infection as a group but not to identify the exact viral pathogen involved.

### Conclusion

In conclusion, a new and simple peripheral blood biomarker DIRECT LS-TA is proposed here which can be readily used in clinical settings. It can be used to differentiate viral infection and inform clinicians on the management of patients. DIRECT LS-TA will emerge as a new kind of in vitro diagnostics (IVD) which can convey single cell-type gene expression information from peripheral blood samples. The new kind of IVD and uniqueness of the information, together with the ease of implementation will make it very useful in clinics.

Antimicrobial resistance remains a global health challenge and accurate and timely discriminating diagnosis of viral infection is essential to reduce antibiotic misuse and overuse (Naghavi et al. 2024). The high correlation of gene expression of these target genes in isolated monocytes and direct measurement of TA in peripheral blood without the need of cell sorting are unique features of DIRECT LS-TA method. This technology is feasible to apply in clinical setting to provide a robust and accurate differential diagnosis of viral infection.

## Acknowledgement

The authors declare the following potential conflict of interest. Nelson LS Tang is the inventor of the patent “Determination of gene expression levels of a cell type “which has been assigned to The Chinese University of Hong Kong. K.S. Leung and Nelson LS Tang are share-holders of Cytomics Ltd. Cytomics Ltd. holds a license to use a patent related to DIRECT LS-TA assay. Patent application pending.

